# Comparative genomics identified a genetic locus in plant-associated *Pseudomonas* spp. that is necessary for induced systemic susceptibility

**DOI:** 10.1101/517870

**Authors:** Polina Beskrovnaya, Ryan A. Melnyk, Zhexian Liu, Yang Liu, Melanie A. Higgins, Yi Song, Katherine Ryan, Cara H. Haney

## Abstract

Plant root-associated microbes promote plant growth and induce systemic resistance (ISR) to foliar pathogens. In an attempt to find novel growth-promoting and ISR-inducing strains, we previously identified strains of root-associated *Pseudomonas* spp. that promote plant growth but unexpectedly induced systemic susceptibility (ISS) rather than ISR to foliar pathogens. Here we demonstrate that the ISS-inducing phenotype is common among root-associated *Pseudomonas* spp. Using comparative genomics, we identified a single *P. fluorescens* locus that is unique to ISS strains. We generated a clean deletion of the 11-gene ISS locus and found that it is necessary for the ISS phenotype. Although the functions of the predicted genes in the locus are not apparent based on similarity to genes of known function, the ISS locus is present in diverse bacteria and a subset of the genes have previously been implicated in pathogenesis in animals. Collectively these data show that a single bacterial locus contributes to modulation of systemic plant immunity.

**Importance:** Microbiome-associated bacteria can have diverse effects on health of their hosts, yet the genetic and molecular basis of these effects have largely remained elusive. This work demonstrates that a novel bacterial locus can modulate systemic plant immunity. Additionally, this work demonstrates that growth promoting strains may have unanticipated consequences on plant immunity and this is critical to consider when engineering the plant microbiome for agronomic improvement.

## Introduction

Plant growth promotion by beneficial microbes has long been of interest because of the potential to improve crop yields. Individual root-associated microbial strains can promote plant growth by facilitating nutrient uptake, producing plant hormones, or improving resilience to both abiotic and biotic stresses (1). In some cases, single bacterial loci underlie beneficial effects of microbes on plants, while other traits appear to be complex and polygenic.

*Pseudomonas fluorescens* and related species are a model for beneficial host-associated microbes due to their genetic tractability and robust host-association across diverse eukaryotic hosts. Direct plant growth promotion (PGP) by *Pseudomonas* spp. can be mediated by bacterial production of the phytohormones auxin (2) or by the expression of 1-aminocyclopropane-1-carboxylate (ACC) deaminase that metabolizes plant-derived ethylene (1, 3). Indirect PGP through antimicrobial activity and pathogen suppression has been attributed to production of the antibiotic 2,4-diacetylphloroglucinol (DAPG) (4). However, the molecular basis of many traits such as induced systemic resistance (ISR) has remained elusive, and multiple distinct bacterial traits including production of siderophores, LPS, and salicylic acid have all been implicated (5).

We previously reported two *Pseudomonas* spp. that induce systemic susceptibility (ISS) on *Arabidopsis* and can promote growth under nutrient limiting conditions (6, 7). These same *Pseudomonas* strains suppress a subset of salicylic acid (SA)-dependent responses and promote resistance to herbivores (7). Although it is possible that ISS-inducing strains contain multiple genetic loci that affect plant growth and pathogen resistance, we hypothesized that a single bacterial trait may be responsible for both the growth and immunity phenotypes of ISS strains.

Growth and immunity have a reciprocal relationship in plants, leading to growth-defense tradeoffs to the extent that plant stunting has been used as a proxy for autoimmunity (8). As a result, we hypothesized that suppression of plant immunity by *Pseudomonas* strains that trigger ISS may be a consequence of PGP activity. The genomes of ISS strains do not contain genes for the ACC deaminase enzyme prevalent in other *Pseudomonas* PGP strains (3); thus, we hypothesized that there may be a distinct mechanism of growth promotion in these strains.

Because of the high density of sampling and genome sequencing within *P. fluorescens* and related species, we reasoned that if ISS is an overlooked consequence of growth promotion then: 1) we should be able to identify additional ISS strains by sampling known PGP strains and additional root-associated strains, and 2) assuming a single unique locus was responsible, that a comparative genomics approach should reveal the underlying genetic basis of ISS.

Here we report that ISS is relatively common among *Pseudomonas* strains within the *P. fluorescens* species complex. We identified new ISS isolates including previously described PGP or environmental isolates and new isolates from *Arabidopsis* roots. Using comparative genomics, we identified a single bacterial locus that is unique to *Pseudomonas* ISS strains. We show that the putative ISS locus is necessary to elicit ISS. While the function of genes in the locus remains elusive, a subset have previously been implicated in pathogenesis, and we found that the locus contributes to rhizosphere growth. Collectively, these data indicate that a single microbial locus contributes to a systemic immune response in a plant host.

## Results

### ISS is a common feature of growth-promoting *Pseudomonas* spp

We previously reported that two strains of *Pseudomonas* (CH229 and CH267) induce systemic susceptibility (ISS) to the foliar pathogen *Pseudomonas syringae* pv. tomato DC3000 (*Pto*) under conditions where a well-characterized ISR strain [*P. simiae* WCS417 (9)] conferred resistance to *Pto* (6, 7). To the best of our knowledge, descriptions of *Pseudomonas*-elicited ISS against bacterial pathogens are limited to *Pseudomonas* sp. CH229 and CH267, strains that were independently isolated from the rhizospheres of wild *Arabidopsis* plants in Massachusetts, USA. We reasoned that if ISS is common among *Arabidopsis*-associated *Pseudomonas* spp., we would be able to identify additional ISS strains from *Arabidopsis* roots from plants growing at distinct sites.

We isolated 25 new fluorescent pseudomonads from wild-growing *Arabidopsis* plants from additional sites in Massachusetts and in Vancouver, Canada. We generated ∼800 bp sequences of a region of the 16S rRNA gene where strains CH229 and CH267 are 99.5% identical, but each shares only <96% identity to the well-characterized ISR strain WCS417. Reasoning that new ISS strains would be closely related to CH267 and CH229, we selected 3 new isolates [1 from Massachusetts (CH235) and 2 from British Columbia (PB101 and PB106)] that were >97% identical to CH267 by 16S rRNA sequence and another 3 (from British Columbia: PB100, PB105 and PB120) that were <97% identical to CH229 and CH267 (Fig. S1). We tested these 6 new rhizosphere *Pseudomonas* isolates for their ability to trigger ISS.

Consistent with the hypothesis that ISS may be common among closely-related PGP *Pseudomonas*, we found that 2 of the 3 strains that were most closely related to CH267 (CH235 and PB101) elicited ISS (Fig. 1). Two strains with <96% identity to CH267 failed to trigger ISS: PB105 triggered ISR and PB100 had no effect on systemic defenses (Fig. 1). PB106 and PB120 consistently enhanced susceptibility in all experiments, but to a more moderate degree (*p<0.1).

**Figure 1.**
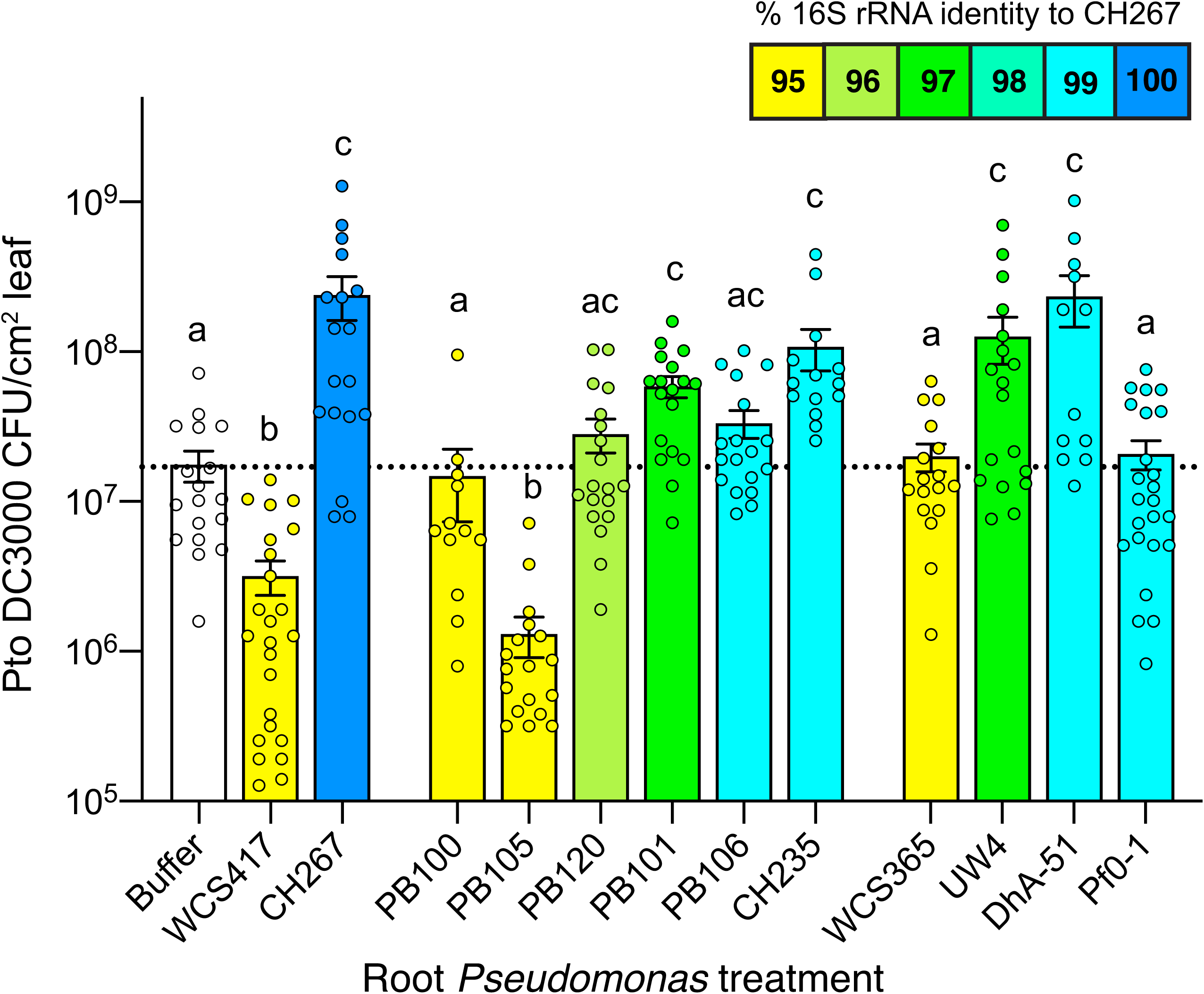
Induced Systemic Susceptibility (ISS) is common among closely-related strains of *Pseudomonas* spp. Isolates of *Pseudomonas* were tested for their ability to modulate systemic defenses; bars are colored to indicate % relatedness to CH267 by partial 16S rRNA sequence as indicated in the key. Data are the average of 3-5 biological replicates with 2 leaves from each of 6 plants (n=12) per experiment. Means +/− SEM are shown. Letters designate levels of significance (p<0.05) by ANOVA and Tukey’s HSD tests.

Collectively, these data indicate that the ability to elicit ISS on *Arabidopsis* ecotype Col-0 may be a common feature among some, but not all, closely-related strains of *Pseudomonas* spp. isolated from the *Arabidopsis* rhizosphere.

Because ISS seemed restricted to strains that were closely related to CH267, we obtained several additional isolates with similar 16S rRNA sequences including *Pseudomonas* sp. UW4, *Pseudomonas* sp. Pf0-1, and *P. vancouverensis* DhA-51. We also tested a growth promoting strain, *Pseudomonas* sp. WCS365 that is more distantly related and to our knowledge has not been tested for ISR/ISS (Table 1). We found that UW4 and DhA-51 elicited ISS while Pf0-1 and WCS365 did not (Fig. 1). *Pseudomonas* sp. UW4 (10) and WCS365 are well-characterized growth promoting strains. *Pseudomonas* sp. Pf0-1 (11) is an environmental isolate. *Pseudomonas vancouverensis* DhA-51 is also an environmental isolate (12) and was previously shown to be closely related to Pf0-1 (13). Because DhA-51 is an environmental isolate that triggers ISS, these data show that the ability to trigger ISS is not specific to rhizosphere isolates.

**Table 1.** Bacterial strains used in this study.

| Strain | Genus and species | Isolated From | Location | Reference |
| --- | --- | --- | --- | --- |
| CH267 | *Pseudomonas* sp. | *Arabidopsis* rhizosphere | Cambridge, MA USA | (6) |
| CH235 | *Pseudomonas* sp. | *Arabidopsis* rhizosphere | Carlisle, MA USA | (6) |
| CH229 | *Pseudomonas* sp. | *Arabidopsis* rhizosphere | Carlisle, MA USA | (6) |
| PB100 | *Pseudomonas* sp. | *Arabidopsis* rhizosphere | Vancouver, BC Canada | This study |
| PB101 | *Pseudomonas* sp. | *Arabidopsis* rhizosphere | Vancouver, BC Canada | This study |
| PB105 | *Pseudomonas* sp. | *Arabidopsis* rhizosphere | Vancouver, BC Canada | This study |
| PB106 | *Pseudomonas* sp. | *Arabidopsis* rhizosphere | Vancouver, BC Canada | This study |
| PB120 | *Pseudomonas* sp. | *Arabidopsis* rhizosphere | Eastham, MA USA | This study |
| WCS417 | *P. simiae* | Wheat rhizosphere | Netherlands | (22) |
| UW4 | *Pseudomonas* sp. | Reeds | Waterloo, ON Canada | (10) |
| Pf0-1 | *Pseudomonas* sp. | Environmental soil |  | (11) |
| DhA-51 | *P. vancouverensis* | Environmental soil | Vancouver, BC Canada | (12) |
| WCS365 | *Pseudomonas* sp. | Tomato rhizosphere | Netherlands | (23) |
| Pf-5 | *Pseudomonas* sp. | Cotton rhizosphere | College Station, TX USA | (24) |
| GW456-L13 | *P. fluorescens* | Groundwater | Oakridge, TN USA | (25) |
| FW300-N1B4 | *P. fluorescens* | Groundwater | Oakridge, TN USA | (25) |
| FW300-N2C3 | *P. fluorescens* | Groundwater | Oakridge, TN USA | (25) |

To gain insights into the distinguishing features of ISS strains, we sequenced the genomes of the 6 new isolates (CH235, PB100, PB101, PB105, PB106 and PB120) from *Arabidopsis* roots as well as *P. vancouverensis* DhA-51 (UW4, WCS365, CH267 and CH229 have been sequenced previously). Whole genome sequencing was used to assemble draft genomes (Methods). We generated a phylogenetic tree using 122 conserved genes as described previously (7, 14). We found that all ISS strains are closely related to one another and fall within a monophyletic group which corresponds to the *P. koreensis, P. jessenii*, and *P. mandelii* subgroups of *P. fluorescens* identified in a recent phylogenomic survey of *Pseudomonas* spp. [Fig. 2B; (15)]. However, not every isolate in this clade is an ISS strain; notably Pf0-1, which has no effect on systemic immunity despite being closely related to CH229. We reasoned that the absence of the ISS phenotype in Pf0-1 should facilitate the use of comparative genomics by allowing us to separate the phylogenetic signature from the phenotypic signature of ISS.

**Figure 2.**
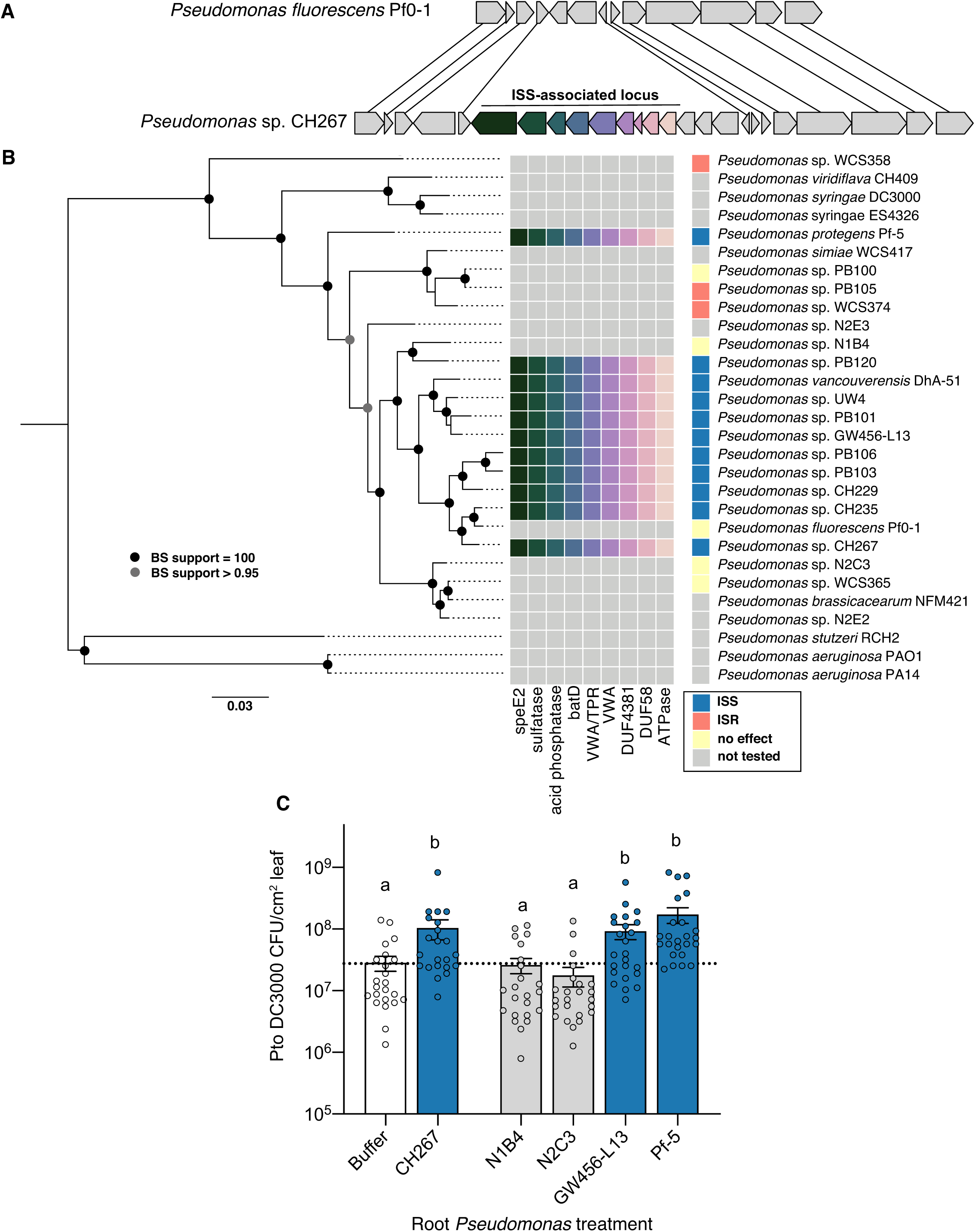
The presence of a genomic island is predictive of the ISS phenotype. **(A)** A genomic island identified through comparative genomics is present in the ISS strains CH229, CH235, CH267 and UW4 and absent in Pf0-1 (no effect on systemic defense) and WCS417 (ISR strain). **(B)** Phylogenetic tree based on 122 core *Pseudomonas* genes. Genome sequencing of new strains shows the island is present in strains that enhance susceptibility but not in those that trigger ISR or have no effect. **(C)** Two strains with the island (GW456-L13 and Pf-5) and two without (N1B4 and N2C3) were tested for ISS/ISR. Only those with the island significantly enhanced susceptibility. Data are the average of 3 biological replicates with 2 leaves from each of 6 plants (n=12) per experiment. Means +/− SEM are shown. *p<0.05 by ANOVA and Tukey’s HSD.

### 11 genes in a single genomic locus are unique to ISS strains and predicts ISS

To identify the potential genetic basis of the ISS phenotype, we used a previously described database of orthologous genes for *Pseudomonas* spp. (14) to identify genes that are present in ISS strains (CH229, CH235, CH267 and UW4) but are absent in the closely-related strain that has no effect on systemic defenses (Pf0-1). We used only the ISS strains with the most robust phenotypes for this analysis. We identified 29 predicted protein-coding genes absent in Pf0-1 but present in all of the other strains. Of these, 12 were small (<100 aa) hypothetical proteins. The remaining 17 predicted protein-coding genes were prioritized for further analysis and are shown in S1 Table. Intriguingly, 11 of the 17 ISS unique genes are found in a single genomic locus.

We surveyed the genomes of other *Pseudomonas* strains tested for ISS to determine if the presence of the 17 genes identified by our comparative genomics approach correlated with the ISS phenotype. We found that the 11 clustered genes were present in ISS strains (DhA-51 and PB101) and the strains with intermediate phenotypes (PB120 and PB106) but were absent in the non-ISS strain WCS365, WCS417 and PB105 (Fig. S2). The remaining 6 genes were all present in WCS365 and/or other non-ISS strains (Fig. S2). We chose to focus on the 11 ISS-unique genes (“ISS locus” hereafter) for further study.

We found that the 11 genes in the ISS locus are found at a single genomic locus in all 4 of the ISS strains (Fig. S3 and Fig. 2A). The flanking regions are conserved in the non-ISS strain Pf0-1 (Fig. 2A), indicating a recent insertion or deletion event. Within this locus, there is a single gene that is conserved in Pf0-1 in addition to two genes that are unique to each individual strain suggesting multiple changes to this genomic region in recent evolutionary history. While all 11 genes are within the same genomic region in the ISS strains, the variability of this locus between closely related strains suggests it may be rapidly evolving.

We surveyed the genomes of sequenced isolates available in our collection for the presence of the ISS locus. We found a number of closely-related strains from various environmental sources that contained the ISS locus, as well as a more distantly related strain (Pf-5) (Fig. 2B). We tested 2 strains that contain the ISS locus (Pf-5 and GW456-L13) as well as 2 that do not (FW300-N1B4 and FW300-N2C3) and found that the presence of the ISS locus correlated with the ISS phenotype, including the distantly-related strain Pf-5 (Fig. 2C). Collectively, these data show that the presence of the 11 candidate genes in the ISS locus identified by our comparative genomics approach is predictive of the ISS phenotype.

### The ISS locus is necessary for ISS

To test if the ISS locus is necessary for ISS strains to induce systemic susceptibility, we deleted the entire 15 kB locus including the region spanning the 11 genes identified in our initial comparative genomics screen in strains CH267 and UW4 (Fig. 2A). We tested these deletion mutants for their ability to induce systemic susceptibility and found that deletion of the entire 11-gene locus (ΔISSlocus), resulted in a loss of the ISS phenotype in both CH267 and UW4 (Fig. 3A and B). This indicates that the ISS locus is necessary for ISS.

**Figure 3.**
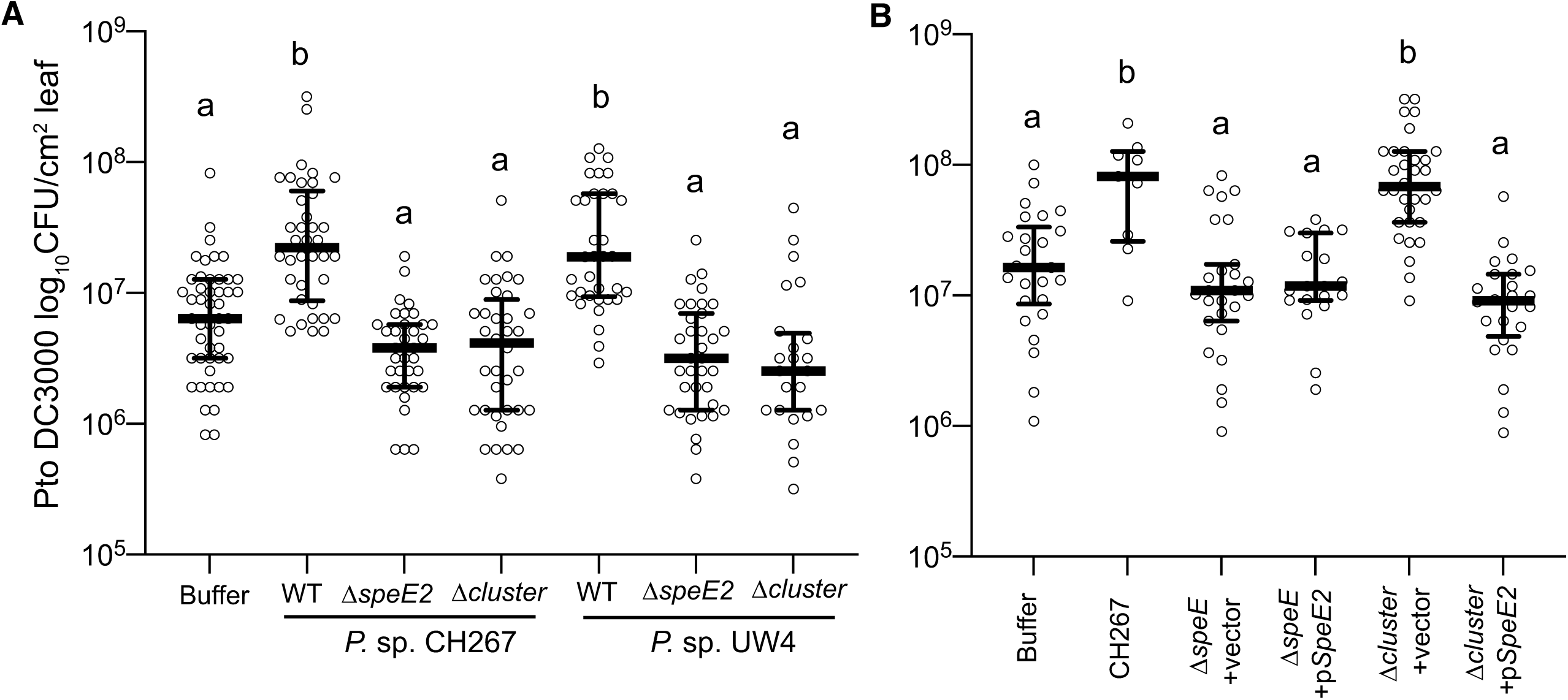
The ISS locus and *speE2* gene are necessary for ISS. **(A-B)** The *speE2* gene and the entire 11-gene locus were deleted from CH267 (A) and UW4 (B). **(C)** Expression of *speE2* from a plasmid is sufficient to complement the CH267 Δ*speE2* mutant but not the ΔISSlocus mutant. Data are the average of 3 biological replicates with 2 leaves from each of 6 plants (n=12) per experiment. Means +/− SEM are shown. *p<0.05 by ANOVA and Tukey’s HSD.

The functions of the majority of the genes in the ISS locus are not apparent based on similarity to genes of known function. A predicted 2544 bp gene was annotated in the CH267 and other genomes as *speE2* due to the similarity of the predicted *C-*terminus to well-characterized spermidine synthase gene *speE1* (PputUW4_02826 and CP336_12795 in UW4 and CH267, respectively). CH267 *speE2* has similarity to a characterized spermidine synthase gene *speE* in *P. aeruginosa* [25% predicted amino acid identity to *P. aeruginosa* PA1687 (16)]. A second *speE*-like gene in the genomes of UW4 and CH267, annotated as *speE1*, is outside of the ISS locus (PputUW4_03691 and CP336_28780 in UW4 and CH267 respectively) and is highly similar to the *P. aeruginosa speE* gene (∼84.0% predicted amino acid identity) (16).

To test if the *speE2* gene is necessary for ISS, we also constructed an in-frame deletion of just the *speE2* gene in both CH267 and UW4. We found that deletion of *speE2* abolished in the ISS phenotype in both CH267 and UW4 (Fig. 2A and 2B) To determine if *speE2* is the only gene within the ISS locus that is necessary for induction of ISS, we generated a complementation plasmid where the CH267 *speE2* gene is expressed under the lac promoter (*p*_*lac*_*-speE2*). We introduced this plasmid into the Δ*speE2* deletion and ΔISSlocus deletions in CH267. While *p*_*lac*_*-speE2* complemented the CH267 Δ*speE2* deletion, it failed to complement the ΔISSlocus deletion (Fig. 3C) indicating that *speE2* is not the only gene within the ISS locus that is required for ISS.

Because deletion of *speE2* in CH267 and UW4 results in the specific loss of the ISS phenotype, this result indicates that the *speE1* and *speE2* genes are not functionally redundant. *SpeE1* and *speE2* differ in length and predicted structure (Fig. 4A). *SpeE1* encodes a predicted 384-amino acid protein and contains a predicted polyamine synthase domain with a predicted decarboxylated S-adenosyl methionine (dSAM) binding motif. *SpeE2* encodes a protein of a predicted 847 amino acids. Similar to *speE1*, the C-terminus of *speE2* contains a predicted dSAM-binding domain; however, *SpeE2* contains predicted transmembrane domains at its N-terminus (Fig. 4A). Spermidine synthases generate spermidine by transferring the aminopropyl group of dSAM to putrescine. Previous structural and mutagenesis analysis on human and *Thermatoga maritima* SpeE1 enzymes revealed common residues important for catalysis (D276, D279, D201, and Y177 for the human SpeE1, and the corresponding D173, D176, D101, and Y76 from the *T. maritima* SpeE1) (17, 18). The catalytic mechanism was proposed to be initiated by the deprotonation of the putrescine amino group by the conserved aspartic acid D276 or D173 with the aid of the side chains of D201 or D101 and Y177 or Y76 as well as the main chain carbonyl of L277 or S174, setting up a nucleophilic attack on dcAdoMet. In addition, residue D279 or D176 is thought to play a role in substrate binding (17, 18).

**Figure 4.**
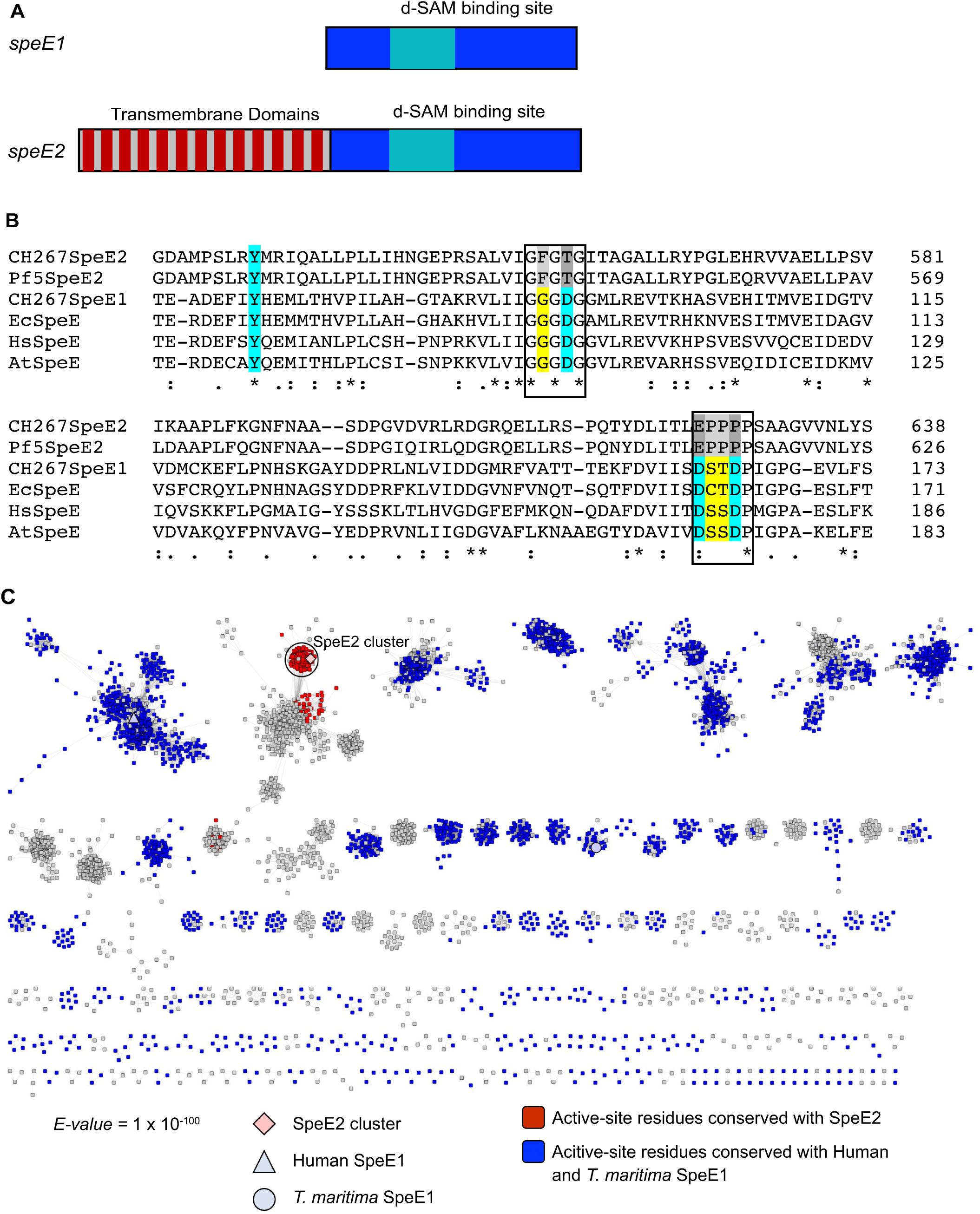
*speE2* is different from characterized spermidine synthases. **(A)** The genome of CH267 contains two *speE* homologues. Both contain predicted d-SAM binding domains and a spermidine synthase domain. Only SpeE2 contains predicted N-terminal transmembrane domains. **(B)** Multiple sequence alignment of predicted amino acid sequence of CH267 SpeE2 and the relatively distantly related Pf-5 SpeE2 gene along with SpeE1-like proteins from CH267, *E. coli, Homo sapiens*, and *Arabidopsis thaliana.* Although the catalytic (blue) and binding-site (yellow) are conserved in all SpeE1 homologues, both SpeE2 genes have changes in these regions (gray). (**C**) Sequence Similarity Network (SSN) of SpeE2 and protein sequences found with the PFAM domain code PF17284. Sequences that have the conserved residues D201/D101, D276/D173, and D279/D176 similar to the human and *T. maritima* SpeE1 are colored blue while sequences that had conserved residues T556, E624, P627 similar to SpeE2 are colored red. Clusters with only 1 sequence were removed for simplicity.

To determine if SpeE2 has the potential to be a spermidine synthase, we performed an amino acid sequence alignment to see if the catalytic residues from classic spermidine synthases are conserved in SpeE2. We found that although the tyrosine residue is conserved, SpeE2 consists of different residues at the corresponding aspartic acid positions. The proposed catalytic residue D276 or D173 in the human or *T. maritima* enzymes corresponds to E624 in SpeE2 while residues D201 or D101 and D279 or D176 have been converted to T556 and P627 (Fig. 4B). Furthermore, we generated a sequence similarity network for SpeE2 with enzymes found in the PF17284 protein family and found that SpeE2 belongs to a distinct cluster away from any functionally characterized enzymes (Fig. 4C). Interestingly, the SpeE2 active site residue substitutions are almost completely conserved within and unique to the SpeE2 cluster (Fig. 4C) suggesting that while *Pseudomonas* sp. CH267 SpeE2 is unlikely to act as a spermidine synthase it may have a distinct function.

### Additional roles for the ISS locus in host interactions

While *speE2* is necessary for ISS, the failure of Δ*speE2* to complement the 11-gene ISS locus deletion (Fig. 3C) indicates that at least one other gene in the ISS locus is likely required for ISS. We tested whether *speE2* is always associated with the same larger locus across the genus *Pseudomonas*. When we analyzed our entire computational dataset of >3800 genomes from across *Pseudomonas*, we found that there was a strong correlation for the presence or absence of 9 of 11 genes (r > 0.9, Fig. 5A). Moreover, we also found that these 9 co-occurring genes were frequently found in the same genomic region, as there were moderate to strong correlations for 9 of the 11 genes co-occurring in the same 50-kb genomic region (Fig. 5B). From a phylogenomic standpoint, we found that these genes were broadly distributed throughout the *Pseudomonas* genus and co-occurred even in taxonomic groups far outside of the *P. fluorescens* clade (Fig. 5C). Within the *P. fluorescens* clade, the ISS locus genes are frequently found in some clades, such as the *koreensis* and *jessenii* clades, which contain most of our isolates (Fig. 5D). However, some clades are missing these genes entirely, such as the plant associated *corrugata* clade (Fig. 5D). Together, these genomic data indicate that despite their polyphyletic distribution among divergent clades of *Pseudomonas* spp., the genes in the ISS locus likely participate in conserved or similar functions.

**Figure 5.**
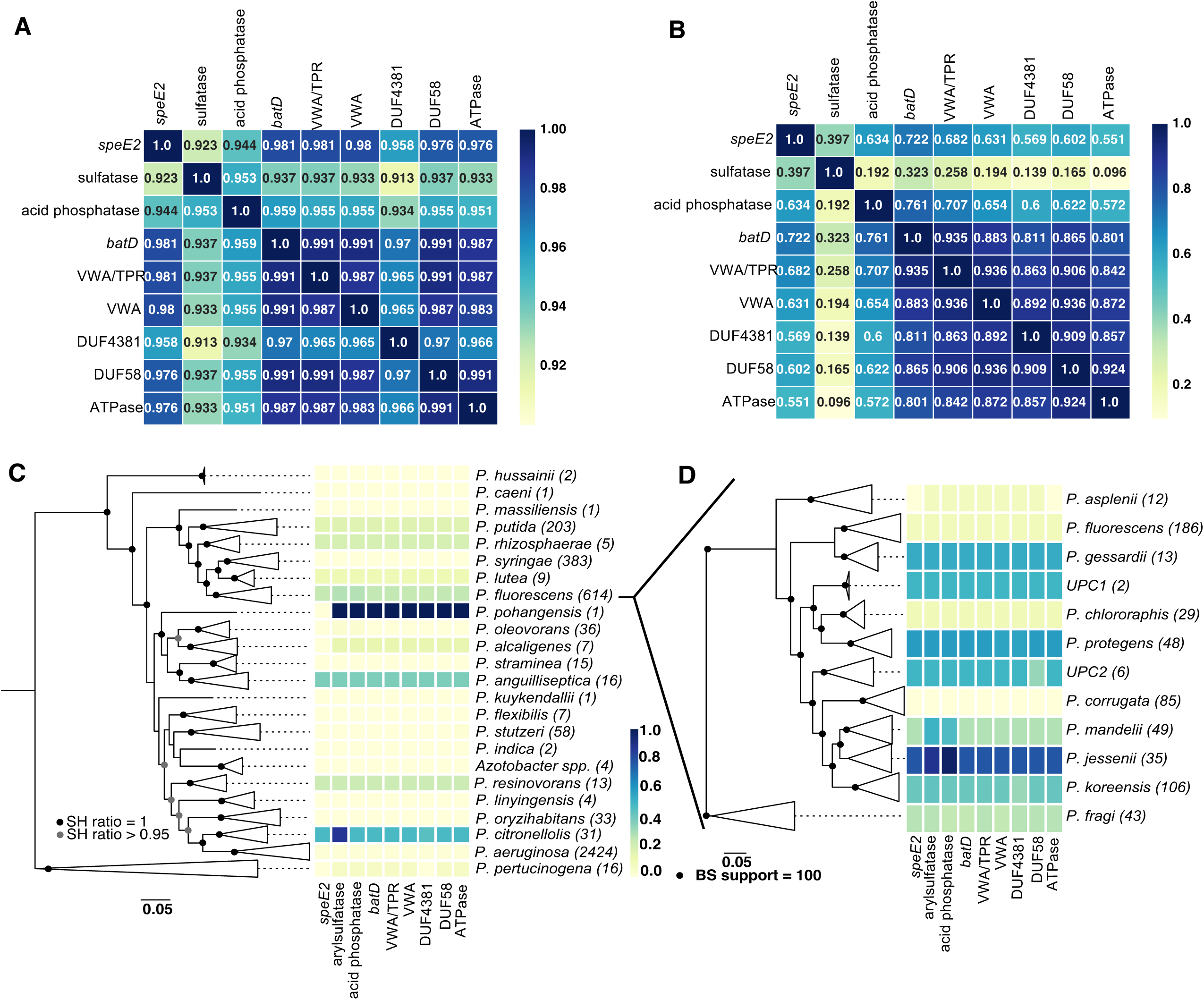
9 genes in the ISS locus nearly always co-occur and are present across the *Pseudomonas* genus. **(A)** Correlation coefficient matrix for 9 genes in the ISS locus across all 3,886 *Pseudomonas* genomes in the comparative genomics database. **(B)** Correlation coefficient matrix for the 9 ISS genes across every 50-kb genomic region that contains at least one of the 9 genes. **(C)** Distribution of the 9 ISS genes across subclades of the *Pseudomonas* genus. (**D**) Distribution of the 9 ISS genes within subclades of the *P. fluorescens* group.

Within the 9 genes that have a high frequency of co-occurrence, we identified a 6 gene predicted operon in the ISS locus with identical domain structure and organization that is involved in stress resistance and virulence in *Francisella tularensis* (19) (Fig. 6A). Another similar operon is associated with aerotolerance and virulence in *Bacteroides fragilis* (20). Returning to our comparative genomics database, we found that these 6 genes comprise an operon broadly conserved in the *Pseudomonas* clade that is distinctly paralogous from the 6-gene operon in the ISS locus (Fig. 6A). This raises the possibility that these six genes within the ISS locus contribute to host-bacterial interactions across diverse bacterial taxa and both plant and animal hosts (Fig. 6A).

**Figure 6.**
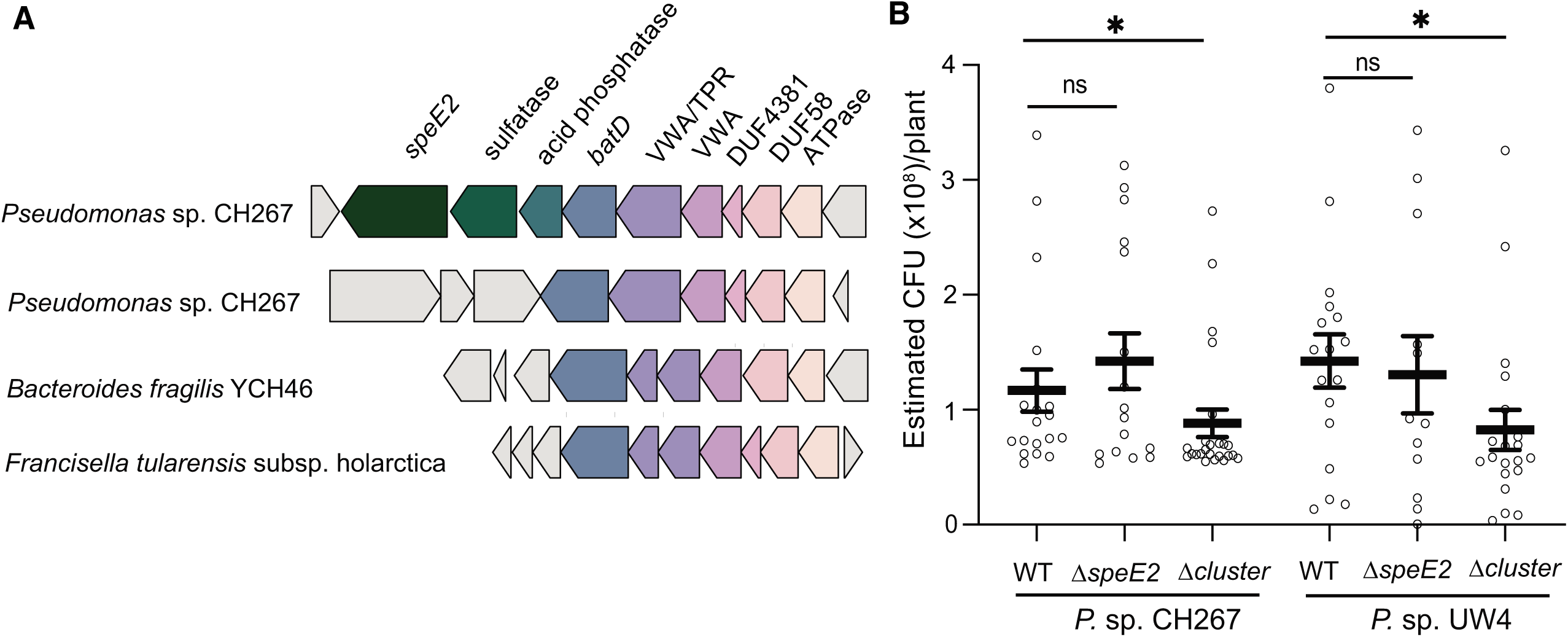
A conserved subset of genes in the ISS locus contribute to virulence and host association in mammalian pathogens and in *Pseudomonas* spp. **(A)** Of the 11 genes in the ISS locus, 6 are contained within a paralogous operon that is present in CH267 and most other *Pseudomonas* spp. An operon with a similar configuration is also present in mammalian pathogens and has been implicated in virulence. (B) The ISS locus, but not the *speE2* gene, promotes rhizosphere colonization. We tested the ΔISSlocus and Δ*speE2* mutant in CH267 and UW4 using a 48-well plate-based rhizosphere colonization assay. Data shown are from 5 days post inoculation. *p<0.05 between mutants in a genetic background by ANOVA and Tukey’s HSD.

To test if the ISS locus is required for *Pseudomonas* to grow in the Arabidopsis rhizosphere, we tested the UW4 and CH267 ΔISSlocus and Δ*speE2* mutants for rhizosphere growth. We transformed the wildtype and mutant CH267 and UW4 strains with a GFP plasmid and used a previously described 48-well plate assay to quantify bacterial growth in the rhizosphere (6). Under these conditions, we observed a significant decrease in rhizosphere growth of ΔISScluster deletion mutants in both the UW4 and CH267 backgrounds (Fig. 6B). We found no decrease in rhizosphere colonization by Δ*speE2* mutants in either the CH267 or UW4 genetic background (Fig. 6B). Together these data indicate that the ISS locus contributes growth in the rhizosphere; however, the Δ*speE2* mutant has a loss of ISS while retaining normal rhizosphere growth indicating a dual role in both rhizosphere colonization and ISS for this genetic locus.

## Discussion

Plant root-associated (“rhizosphere”) microbes perform a diversity of functions that benefit their plant hosts including nutrient uptake and defense. Functional characterization of individual plant-associated bacterial and fungal strains of potential agronomic importance (i.e. growth promoters or nitrogen fixers) is widespread (5). However, closely-related strains of bacteria can have very distinct effects on plant growth and defense (13), and these effects can be dependent on environmental context (1). Lack of known correlations between microbial genotype and potential effects on plant hosts present a challenge when attempting to infer the effect that a microbe may have on its plant host from sequence identity alone.

Our use of comparative genomics and isolate phenotyping to identify the genetic basis of a complex microbial-derived trait indicates that this is an effective approach to identifying important microbial traits to improve plant health. For comparative genomics to be effective, traits should be controlled by single or limited genomic loci, and phylogeny should not be predictive of function. In this case, a close relative of ISS strains, *Pseudomonas* sp. Pf0-1 (>99% identical by full length 16S rRNA to the ISS strains) does not affect systemic defenses (Fig. 1), which allowed us to use comparative genomic to identify the underlying basis. We previously used this approach to find the genomic basis of a pathogenic phenotype within a clade of commensals (14). It has been previously observed that phylogeny is not predictive of function for ISR strains (13) suggesting that comparative genomics may be appropriate to find the basis of additional plant-associated traits.

We found that the ISS locus encodes genes involved in both triggering ISS and promoting rhizosphere colonization. Loss of the entire locus results in a loss of ISS and a decrease in growth in the rhizosphere; however, loss of *speE2* impairs ISS but not rhizosphere growth suggesting that there may be multiple plant-association functions encoded in this locus. The function of the *speE2* gene and other genes within the ISS locus are not readily apparent from similarity to previously characterized enzymes. As spermidine and other polyamines should directly enhance plant resistance through generation of ROS (21), it is possible that the *speE2* enzyme converts spermidine or another polyamine to a non-defense inducing molecule. The highly conserved nature of the active-site residues within *speE2-*like genes suggests a novel function in this enzyme.

While enhancement of systemic susceptibility is not an obviously agronomically useful plant trait, several ISS strains promote growth and enhance resistance to insect pests (6, 7). Using ISS strains might be beneficial for crops where insects are the primary pressure on crop productivity. However, the ubiquity of ISS by plant growth-promoting strains illustrates the complexity of host-microbe interactions and should be considered when engineering the microbiome.

## Materials and Methods

### Plant growth conditions

For all experiments, plants were grown in Jiffy-7 peat pellets (Jiffy Products) under a 12 h light/12 h dark at 22 °C temperature regime. Seeds were surface sterilized by washing with 70% ethanol for 2 minutes followed by 5 minutes in 10% bleach and 3 washes in sterile water. Seeds were stored at 4° C until use. Unless otherwise indicated, seeds were sowed in Peat pellets (Jiffy 7) and placed in a growth chamber under 12-hour days and 75 μM cool white fluorescent lights at 23° C.

### Bacterial growth and 16S rRNA sequencing

*Pseudomonas* strains were cultured in LB or King’s B at 28 °C. New *Pseudomonas* strains were isolated from the roots of wild-grown *Arabidopsis* plants around eastern Massachusetts, USA and British Columbia, Canada as described (6). New *Pseudomonas* isolates were preliminary identified based on fluorescence on King’s B and confirmed by 16S rRNA sequencing.

### ISS assays

ISS and ISR assays were performed as described (7, 26). Briefly, *Pseudomonas* rhizosphere isolates were grown at 28 °C in LB medium. For inoculation of plant roots for ISR and ISS assays, overnight cultures were pelleted, washed with 10 mM MgSO_4_ and resuspended to a final OD_600_ of 0.02. Jiffy pellets were inoculated 9 days after seed germination with 2 mls of the indicated bacterial strains at a final OD_600_ of 0.02 (5×10^5^ CFU g^−1^ Jiffy pellet). For infections, the leaves of 5-week old plants were infiltrated with *Pto* DC3000 at an OD_600_ = 0.0002 (starting inoculum ∼10^3^ CFU/cm^2^ leaf tissue). Plants were maintained under low light (<75 µM) and high humidity for 48 hours. Leaf punches were harvested, ground, and plated to determine CFU counts.

### 16S rRNA sequencing, bacterial genome sequencing, assembly and phylogenomics

Bacterial DNA preps were performed using Qiagen Purgene Kit A. 16S rRNA was amplified using 8F and 1391R and sequenced using 907R. Bacterial genomic library prep and genome sequence was performed as described (7). Briefly, bacterial DNA was isolated using Qiagen Purgene Kit A and sonicated into ∼500 bp fragments. Library construction was performed as described (7), individually indexed and sequenced using MiSeq V3 paired end 300 bp reads. After barcode splitting, approximately 500,000 to 1 million reads were used for each sample to assemble draft genomes of the strains *Pseudomonas* sp. CH235, PB100, PB101, PB105, PB106, PB120 and *P. vancouverensis* DhA-51. Genome assembly was carried out as previously described (7) and draft genomes are available from NCBI (see below).

### Phylogenomic tree building

To generate the 29-taxon species tree used in Figs. 2B and 4E, we made use of an alignment of 122 single-copy genes we previously found to be conserved in all *Pseudomonas* strains (14). From this amino acid alignment, we extracted 40,000 positions ignoring sites where >20% of the taxa had gaps. Using RAxMLv8.2.9, we inferred 20 independent trees under the JTT substitution model using empirical amino acid frequencies and selected the one with the highest likelihood. Support values were calculated through 100 independent bootstrap replicates under the same parameters.

To build the 3,886-taxon phylogeny of the *Pseudomonas* genus in Figs. 5C and S1, the same 122-gene alignment was used. For computational feasibility, the alignment was randomly subsampled to 10,000 amino acid positions, again ignoring sites that were highly gapped (>20%). FastTree v2.1.9 was used to build the phylogeny using default parameters. The phylogeny was rooted to a clade of *Pseudomonas* identified as an outgroup to all other *Pseudomonas* spp. as previously described (14). To more easily visualize this tree, we collapsed monophyletic clades with strong support (as determined by FastTree’s local Shimodaira-Hasegawa test) that correspond with major taxonomic divisions identified by Hesse et al. (2018).

To build the tree for the *Pseudomonas fluorescens* (*Pfl*) subclade seen in Figs. 5D and S2, we identified 1,873 orthologs specific to the *Pfl* clade found in >99% of all strains in the clade and then aligned them all to the hidden Markov models generated by PyParanoid using hmmalign, prior to concatenation. This alignment had 581,023 amino acid positions, which we trimmed to 575,629 positions after masking sites with >10% of taxa with gaps. From this alignment, we randomly subsampled 120,000 sites for our final phylogenomic dataset. Using RAxMLv8.2.9, we inferred 20 independent trees under the JTT substitution model using empirical amino acid frequencies and selected the one with the highest likelihood. Support values were calculated through 100 independent bootstrap replicates under the same parameters.

### Comparative Genomics

Comparative genomics analyses were performed by using a previously described framework for identifying PyParanoid pipeline and the database we built for over 3800 genomes of *Pseudomonas* spp. Briefly, we had previously used PyParanoid to identify 24,066 discrete groups of homologous proteins which covered >94% of the genes in the original database. Using these homolog groups, we annotated each protein-coding sequence in the newly sequenced and merged the resulting data with the existing database, generating presence-absence data for each of the 24,066 groups for 3,886 total *Pseudomonas* genomes.

To identify the groups associated with induction of systemic susceptibility, we compared the presence-absence data for 4 strains with ISS activity (*Pseudomonas* spp. CH229, CH235, CH267, and UW-4) and 1 strain with no activity (*Pseudomonas* sp. Pf0-1). We initially suspected that ISS activity was due to the presence of a gene or pathway (i.e. not the absence of a gene) and thus initially focused on genes present only in Pf0-1. We identified 29 groups that were present in the 4 ISS strains but not in Pf0-1.

To obtain the correlation coefficients in Figs. 4D and 5A, we coded group presence or absence as a binary variable and calculated Pearson coefficients across all 3,886 genomes. To calculate the correlation coefficients in Fig. 5B, we split the genomic database into 50-kb contiguous regions and assessed group presence or absence within each region. Because this dataset is heavily zero-inflated, we ignored regions that had none of the 11 groups, taking the Pearson coefficient of the 11 genes over the remaining regions.

Initial annotation of the ISS groups was based on generic annotations from GenBank Further annotation of the 11 groups specific to the ISS locus was carried out using the TMHMM v2.0 server, the SignalP 4.1 server and a local Pfam search using the Pfam-A database from Pfam v31.0. To identify homologous genes in the genomes of *Francisella tularensis* subsp. *holarctica* and *Bacteroides fragilis* YCH46, we relied on locus tags reported in the literature which we confirmed using annotation based on another Pfam-A domain search.

### Deletion of the *speE2* gene and 11-gene ISS locus

Deletions in the CH267 and UW4 strains were constructed by a two-step allelic exchange as described (27). The flanking regions directly upstream and downstream of the 11-gene ISS locus or the *speE2* gene were amplified and joined by overlapping polymerase chain reaction (PCR) using genomic DNA as template and primers listed in Table 2. Following digest, the product was ligated into the pEXG2 suicide vector that contains the *sacB* gene for counter-selection on sucrose (28). The recombinant plasmid was then transformed into calcium-competent *E. coli* DH5α by heat shock. After confirmation of correct insert by PCR and sequencing, the plasmid was transformed into WM3064 (29). Conjugation of plasmid into CH267 and UW4 from WM3064 was performed by biparental mating on King’s B media supplemented with diaminopimelic acid, and transconjugants were selected using 10 µg/mL gentamicin and 15 µg/mL nalidixic acid. The second recombination leading to plasmid and target DNA excision was selected for by using sucrose counter-selection. Gene deletions in CH267 and UW4 were confirmed by PCR amplification of the flanking regions with primers listed in Table 2, agarose gel electrophoresis and Sanger sequencing.

**Table 2.**
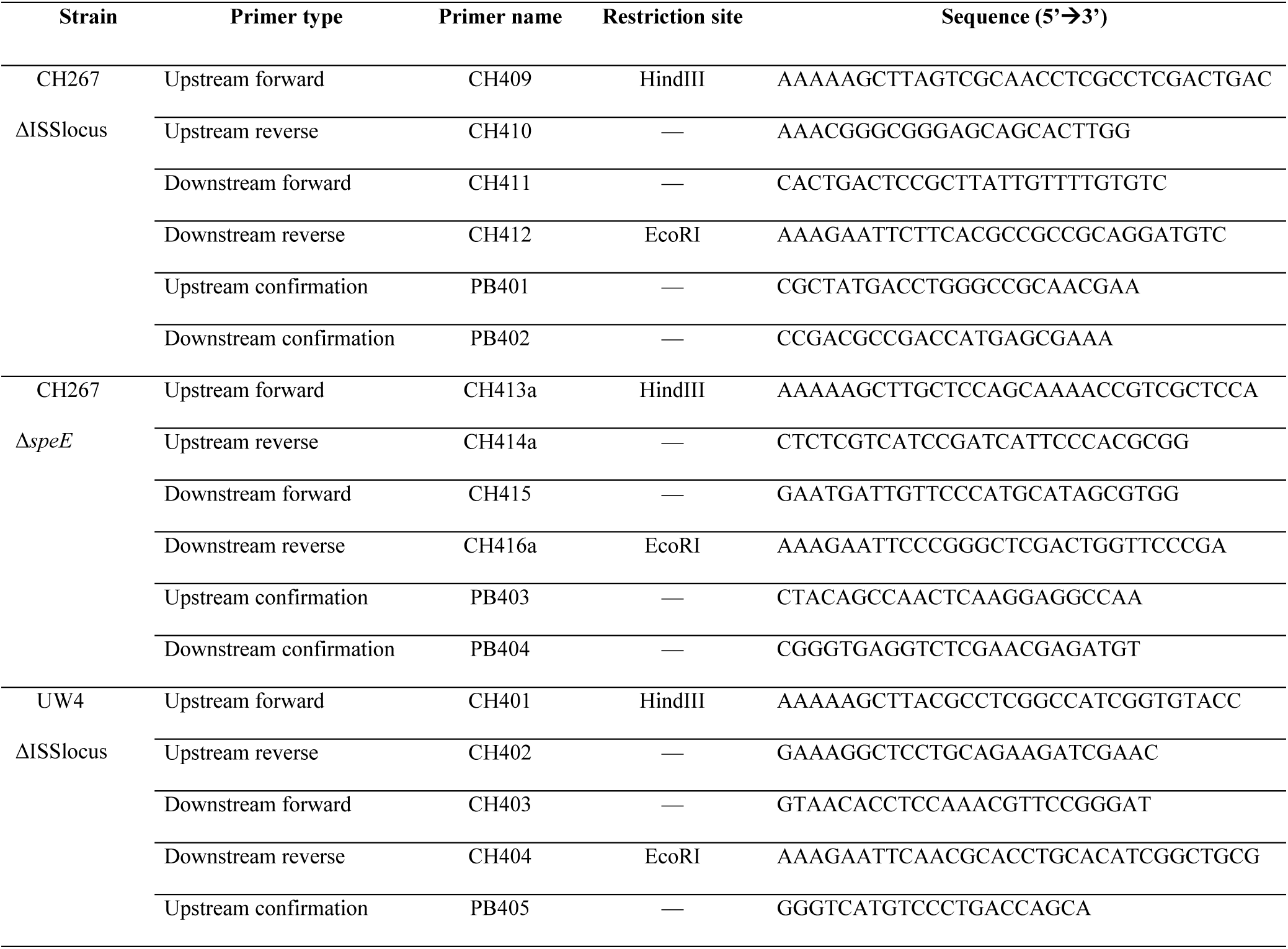
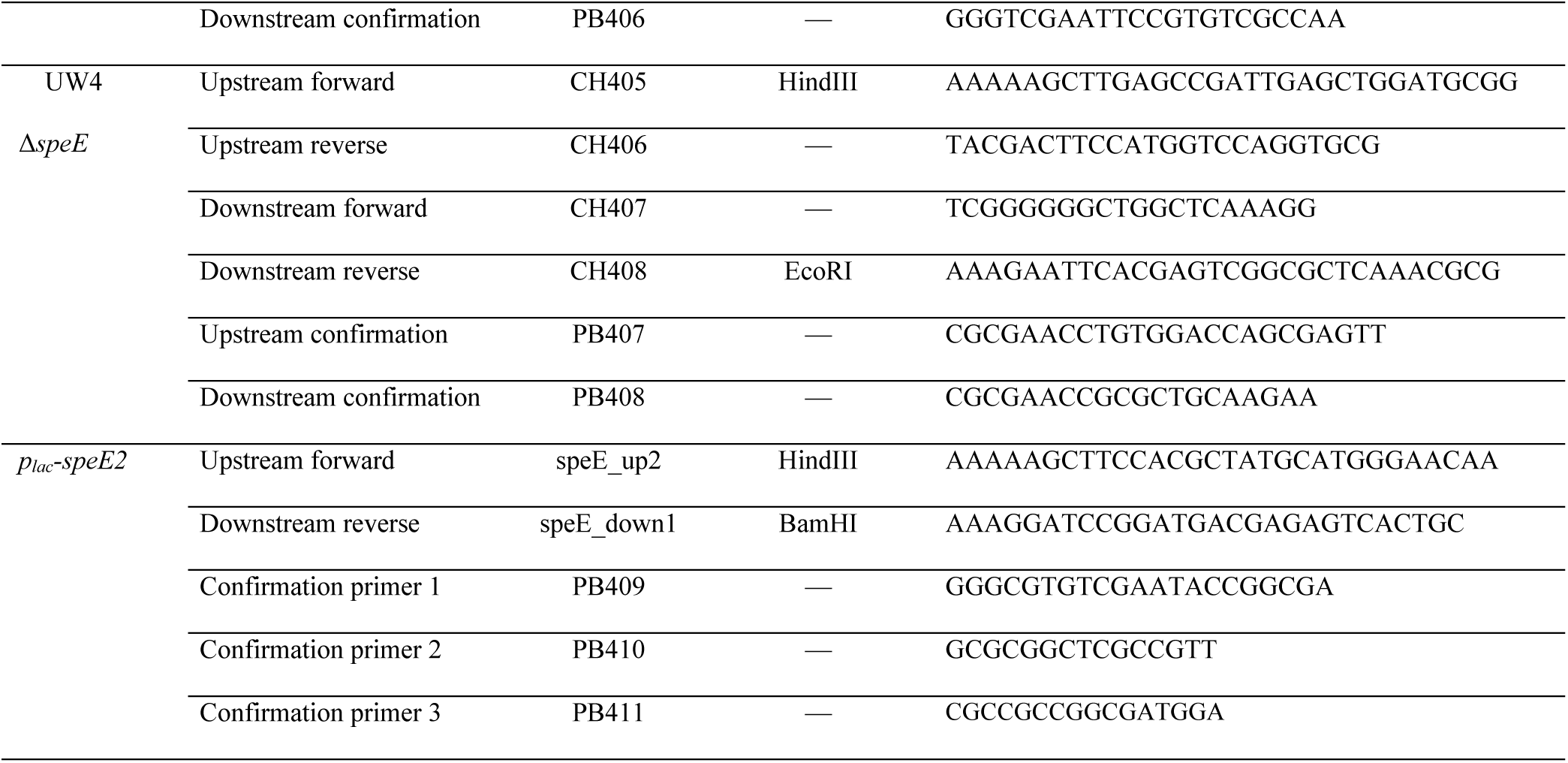
Primers used to generate the mutant *Pseudomonas* strains analyzed in this study.

### Complementation of the *speE2* gene

The *speE2* gene was amplified by PCR using CH267 genomic DNA as template, as well as the primers listed in Table 2. Following restriction digestion, the ∼2.6 kb insert was ligated into the pBBR1MCS-2 vector into the multiple cloning site located downstream of a lac promoter. Ligation mixture was then introduced into *E. coli* DH5α by heat shock, and transformants were selected using LB media supplemented with 25-50 µg/mL kanamycin. Presence of correct insert was confirmed by PCR, restriction digest and Sanger sequencing. pBBR1-MCS2::p_lacZ_-speE2_CDS_ plasmids were maintained in *E. coli* DH5α λpir with 25 μg/mL of Kanamycin. To construct a conjugating strain, Calcium-competent *E. coli* WM3064 was first transformed with pBBR1-MCS2::p_lacZ_-speE2_CDS_ or pBBR1-MCS2 by heat shock. To conjugate *Pseudomonas* sp. CH267, 1 mL of overnight cultures of *Pseudomonas* sp. CH267 and *E. coli* WM3064 carrying the appropriate plasmids were washed twice and resuspended with 0.5 mL of 100 mM MgCl_2_. The resuspended *Pseudomonas* sp. CH267 was mixed with *E. coli* WM3064 strains at 1:2 ratio. Six 25 μL mating spots were placed on LB plates supplemented with 0.3 mM of Diaminopimelic acid (DAP). The mating spots were allowed to dry before incubating at 28°C for 4 hr. The mating spots were then scraped off and resuspended in 1 mL of 100 mM MgCl_2_. 100 μL of the suspension was plated on LB-Kanamycin. Colonies were restreaked to confirm antibiotic resistance.

### Multiple sequence alignment and Sequence Similarity Network (SSN) Generation

Multiple sequence alignment was performed with Clustal Omega (30). The SSN was created using the enzyme function initiative (EFI-EST) web tool (31) by inputting the SpeE2 amino acid sequence with the amino acid sequences from the spermidine synthase tetramerization domain with the code PF17284 using UniRef90 seed sequences instead of the whole family. Sequences will less than 100 amino acids were also excluded resulting in a total of 6523 sequences. An alignment score threshold or E-value cutoff of 10^−100^ was used to generate the SSN which was visualized using Cytoscape (32).

### Rhizosphere colonization assay

Arabidopsis seedlings were grown in 48-well plates and rhizosphere growth of bacteria was quantified as previously described (6). Briefly, *Arabidopsis* seeds were placed individually in 48-well clear-bottom plates with the roots submerged in hydroponic media (300 µl 0.5× MS media plus 2% sucrose). The medium was replaced with 270 µl 0.5× MS media with no sucrose on day 10, and plants were inoculated with 30 µl bacteria at an OD_600_ of 0.0002 (final OD_600_, 0.00002; ∼1,000 cells per well) on day 12. Plants were inoculated with wild-type *Pseudomonas* CH267 or UW4 strains containing plasmid pSMC21 (*pTac-GFP*) (33). Fluorescence was measured with a SpectraMax i3x fluorescence plate reader (Molecular Devices) (481/515 excitation/emission) 5 days post inoculation. A standard curve related fluorescence to OD was generated to estimate CFU/wells (OD_600_ = 1 = 5 × 10^8^ CFU/mL).

## Acknowledgements

This work was supported by an NSERC Discovery Grant (NSERC-RGPIN-2016-04121) and a Seeding Food Innovation grant from George Weston Ltd. awarded to C.H.H. Additional support from a Life Sciences Research Foundation Fellowship from the Simons Foundation awarded to R.A.M., a fellowship from China Postdoctoral Science Foundation awarded to Y.S., a Chinese Graduate Scholarship Council Award to Y.L., and an NSERC CGS-M award to Z. L.

## Author Contributions

C.H., R.A.M., and P.B. designed experiments. P.B. Y.S. Y.L. and C.H.H. performed experiments. C.H., R.A.M., Z.L. analyzed data and R.A.M. performed genome assembly, annotation, phylogenetic analysis and comparative genomics. M.H. and K.R. performed bioinformatic analyses of *speE2* function. C.H.H., P.B. and R.A.M. wrote the manuscript with input from all.

## Data Availability

Data for the Whole Genome Shotgun project has been deposited at DDBJ/ENA/GenBank under the accessions RRZJ00000000 (CH235), RRZK00000000 (DhA-51), RWIM00000000 (PB106), RWIN00000000 (PB120), RWIO0000000 (PB105), RWIQ00000000 (PB100), and RWIR00000000 (PB101). The versions described in this paper are versions RRZJ01000000 (CH235), RRZK01000000 (DhA-51), RWIM01000000 (PB106), RWIN01000000 (PB120), RWIO0100000 (PB105), RWIQ01000000 (PB100), and RWIR01000000 (PB101).

## Declaration of interests

The authors declare no competing interests.

---

**S1 Table. Unique loci identified in comparative genomics.** The genome content of 4 ISS strains (CH267, CH235, UW4 and CH229) was compared with the closely-related non-ISS strain Pf0-1. 17 predicted protein-coding genes were identified.

**Figure S1. Correlation matrix of 16S rRNA similarity of new *Pseudomonas* isolates from the Arabidopsis rhizosphere.** Isolates were selected based on similarity (>97% identical by partial 16S rRNA) to CH267 (CH235, PB101 and PB106) or distance (<97% identity by partial 16S rRNA) to CH267 (PB120, PB100, PB105). Isolates from the rhizosphere of Arabidopsis growing in *Massachusetts, USA or #British Columbia, Canada.

**Figure S2. Distribution of loci identified by comparative genomics ISS loci across *Pseudomonas* strains.** Comparative genomics between ISS strains UW4, CH229, CH235 and CH267 (black arrows) and non-ISS strain Pf0-1 (red arrow) identified 17 predicted protein-coding genes >100 aa that were absent in Pf0-1 and present in strains that induce ISS. 11 of these genes were found in a single genomic locus (box) and were absent in the non-ISS strain WCS365.

**Figure S3. The ISS locus is highly variable between closely-related strains**

The 11 genes in the ISS locus are present in the ISS strains Pf0-1, CH235, CH267 and CH299 but absent in Pf0-1. Genes in the ISS locus are colored as in the key at the bottom of the figure and in Fig. 2. Conserved genes not unique to the ISS strains are colored similarly among strains; genes in gray are not conserved between strains at this locus. In CH229, Pf0-1 and CH267 the genes flanking the ISS locus are conserved in the same orientation suggesting a recent insertion or deletion event.

## Notes

### Competing Interest Statement

The authors have declared no competing interest.

